# Minute virus of mice is oncolytic for pancreatic cancer cells with mesenchymal phenotype

**DOI:** 10.1101/2023.04.11.536425

**Authors:** P. Garcin, H. Lulka, N. Dusetti, M. Vienne, L. Buscail, P. Cordelier

## Abstract

Pancreatic cancer will soon become the second cause of death by cancer in Western countries. The main barrier to increase the survival of patients with this disease requires the development of novel and efficient therapeutic strategies, that better consider tumor biology. Oncolytic viruses are quickly moving toward the forefront of medicines for the management of cancer. Among them, the fibrotropic minute virus of mice prototype (MVMp) preferentially infects migrating and undifferentiated cells, that highly resemble poorly differentiated, basal-like, pancreatic tumors showing the worst clinical outcome. Thus, we hypothesized that MVMp may specifically target the most aggressive subtype of pancreatic cancer cells. We report here that MVMp specifically infects, replicates in and kills pancreatic cancer cells from murine and human origin with a mesenchymal, basal-like profile, while sparing cancer cells with an epithelial phenotype. *In vivo*, MVMp shows oncolytic activity in experimental tumors with mesenchymal phenotype only, and shows increased antitumoral efficacy in immune competent models. Collectively, we demonstrate herein for the first time that MVMp is specific and oncolytic for pancreatic tumors with mesenchymal, basal-like profile, paving the way for precision medicine opportunities for the management of the most aggressive and lethal form of this disease.

## INTRODUCTION

Pancreatic ductal adenocarcinoma (PDAC) is the most common form of pancreatic cancer and has often been viewed as a disease with a poor prognosis^1^. Over the last decade, multiple groups have characterized the complex molecular landscape of PDAC to reveal several distinct classes of disease^2^. Two main molecular classification emerged, including a classical subtype and a basal-like gene expression program that resemble the basal subtype of breast and bladder cancers. with the basal-like subtype showing the poorer clinical outcome^3–5^. However, the question remains on how the recent progress in molecular stratification may translate into tangible benefit for patient with PDAC. With this in mind, we asked whether clinically relevant PDAC molecular subtypes may respond differently to innovative therapies such as oncolytic virotherapy.

Oncolytic virus (OV) exploits the frequent defects in antiviral signaling in cancer cells to prosper and kill cancer cells, leaving normal cells unharmed^6^. Virotherapy is now recognized as a promising multi-mechanistic therapy for the treatment of cancer, with the first OV product Talimogene laherparepvec (T-Vec, Imlygic®) approved by the U.S. FDA in 2015 for the treatment of advanced melanoma^7, 8^. OVs come into different sizes and flavors; among them, the Minute Virus of Mice prototype (MVMp) is a non-enveloped, small single-stranded DNA virus with an icosahedral capsid of 25nm^9^. MVMp is member of the *Protoparvovirus* genus, is nonpathogenic in humans and has potential utility as cancer therapeutic in experimental models of glioma^10^.

MVMp tropism for cancer cells is still not well understood. The role of type-I interferons in limiting MVMp infection is highly debated^11, 12^; interestingly, as antiviral signaling is commonly silenced in PDAC cells^13^, this may provide favorable ground for MVMp replication. Recent works demonstrate the role of sialic acid structures^14^ and galectin-3^15^ in MVMp cell surface binding and entry. Remarkably, MVMp infection is promoted by epithelial-mesenchymal transition (EMT)^16^, a situation that closely resemble basal-like phenotype of PDAC tumors, as MVMp preferably enters leading edge of migrating cells by endocytosis close to filopodial and focal adhesions areas^16^. During this work, we asked whether PDAC cells with basal-like, mesenchymal phenotype might be target for MVMp infection. We identified a 5-gene classifier (FAK, N-cadherin, E-cadherin and β-catenin) that predicts MVMp cytotoxic activity in PDAC cells from murine and human origin with mesenchymal profile, when cells with a more epithelial phenotype were spared by the virus. MVMp tropism was maintained *in vivo*, as MVMp replicated in, inhibited the progression and induced cell death by apoptosis of PDAC experimental tumors with mesenchymal phenotype only. Last, we found that MVMp partners with the immune system to inhibit PDAC growth, with evidence of adaptive immunity engagement. Collectively, we demonstrate here for the first time that MVMp is oncolytic for the most aggressive form of PDAC, paving the way for new precision medicine opportunities for the therapy of this disease with no cure.

## RESULTS

### Isolation of mesenchymal and epithelial subtypes of pancreatic cancer cells

Recent studies demonstrate that basal-like and classical cell types coexist within PDAC^17^. To interrogate how these subtypes respond to virotherapy, we generated long-term cultures of murine R211 and R259 PDAC cells that derive from KPC spontaneous mouse model of cancer (Pdx1-Cre, lox-stop-lox-KrasG12D/+, lox-stop-lox-tp53R172H/+)^18^. We identified clusters of differentiated cells with an epithelial phenotype coexisting with fibroblast-like, less differentiated cells, highly resembling human tumors (Figure 1A and FigS1A). Cells from each phenotype were separated by differential adhesion to plastic, and thereafter named R211-M and R259-M for mesenchymal, and R211-E and R259-E for epithelial (Figure 1A and FigS1A). We then perform western blot analysis for markers of EMT (FAK, N-cadherin, E-cadherin, β-catenin, Slug, Snail) and used β-actin and GAPDH as loading controls. Molecular exploration demonstrates that mesenchymal markers, such as FAK, N-cadherin and Snail, are expressed at high levels in R211-M as compared to R211-E cells (Figure 1B). Conversely, epithelial markers such as E-cadherin and β-catenin are absent in cells with the mesenchymal phenotype, while expressed at high levels in cells with the epithelial phenotype (Figure 1B). We next addressed the ability of these cultures to form tumors in immune deficient mice. As shown in Figure 1C and 1D, engraftment in the pancreas of both R211-M and R211-E led to robust tumor formation, confirming the neoplastic nature of these cultures, with R211-M tumors growing faster than R211-E tumors. Macroscopic analysis of tumors at endpoint reveals that both phenotypes are conserved within experimental tumors, with R211-M and R259-M cells giving rise to poorly differentiated PDAC with extensive desmoplastic stroma (Figure 1E and FigS1C), and R211-E and R259-E cells at the origin of moderately differentiated PDAC embedded in desmoplastic stroma (Figure 1C and FigS1C).

**Figure 1.**
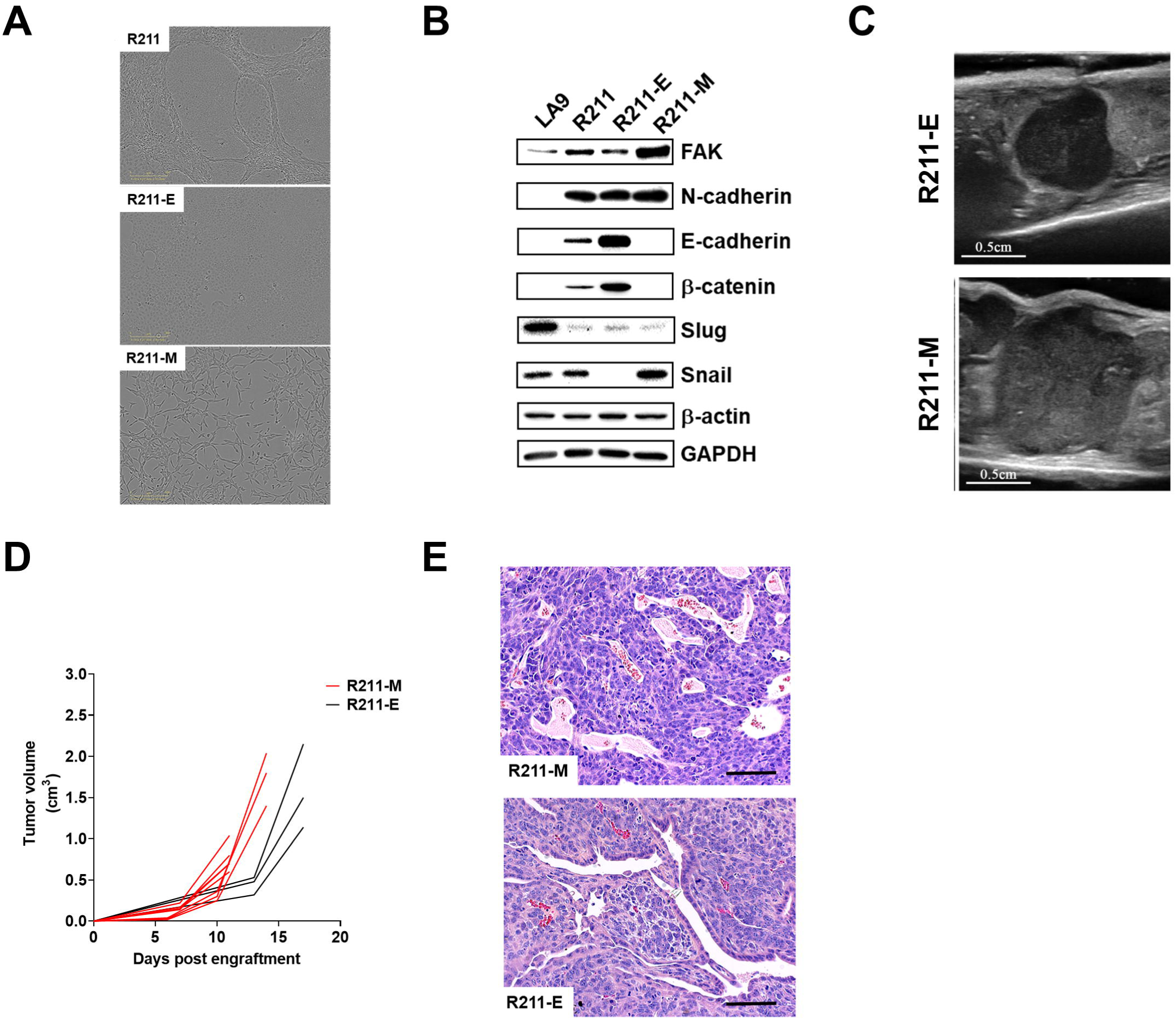
Isolation of mesenchymal and epithelial subtypes of murine pancreatic cancer cells. R211-E and R211-M cells were isolated by differential adherence to plastic as mentioned in the Materials and Methods section. (**A**) Representative captions of long term culture of R211 cells (top) and of isolated R211-E (middle) and R211-M (bottom) cells. **B.** Western blot analysis for the indicated proteins in LA9, parental R211, R211-E and R211-M cells. Representative of two independent experiments. Representative captions of tumor echography (**C**) and tumor growth kinetics (**D**) analyzed by echography of R211-M and R211-E tumors engrafted in NSG mice. N=3 to 8 mice per group. **E.** hematoxylin and eosin (HE) staining of orthotopic R211-E and R211-M tumors engrafted in nude mice at endpoint. Representative of n=3 mice per group.

### Murine pancreatic cancer cells with mesenchymal phenotype are targets for MVMp infection

We next addressed the permissiveness of R211-M and R211-E cells to MVMp infection. For comparison, LA9 cells, the prototypical cellular model for the study for MVMp infection, were used as control. Oncolytic cell death was monitored in real-time using the cytotox green dye and the Incucyte Zoom (FigS2A). As expected, LA9 cells were highly sensitive to MVMp oncolytic effect (Figure 2A and FigS2A), with more than 20-fold increase in cell death as compared to control, 72h following infection (Figure 2B). Remarkably, MVMp infection induced strong cell killing of R211-M cells at MOI=100 (8-fold increase, p<0.001, Figure 2A and Figure 2B), while R211-E cells largely resisted viral-induced oncolysis when challenged with a similar dose of virus. Western blot analysis for PARP cleavage indicate that MVMp infection induces cell death by apoptosis in R211-M cells (Figure 2C), but not in R211-E cells (FigS2B). Plaque assay confirmed that R211-M cells only are sensitive to MVMp infection, albeit at lower level as compared to LA9 cells (FigS2C). We also found that R259 with mesenchymal phenotype were killed by the virus, when cells with epithelial phenotype resisted MVMp infection (FigS2D-E). Thus, we demonstrate that infection by MVMp specifically kills murine PDAC cells with a mesenchymal phenotype *in vitro*.

**Figure 2.**
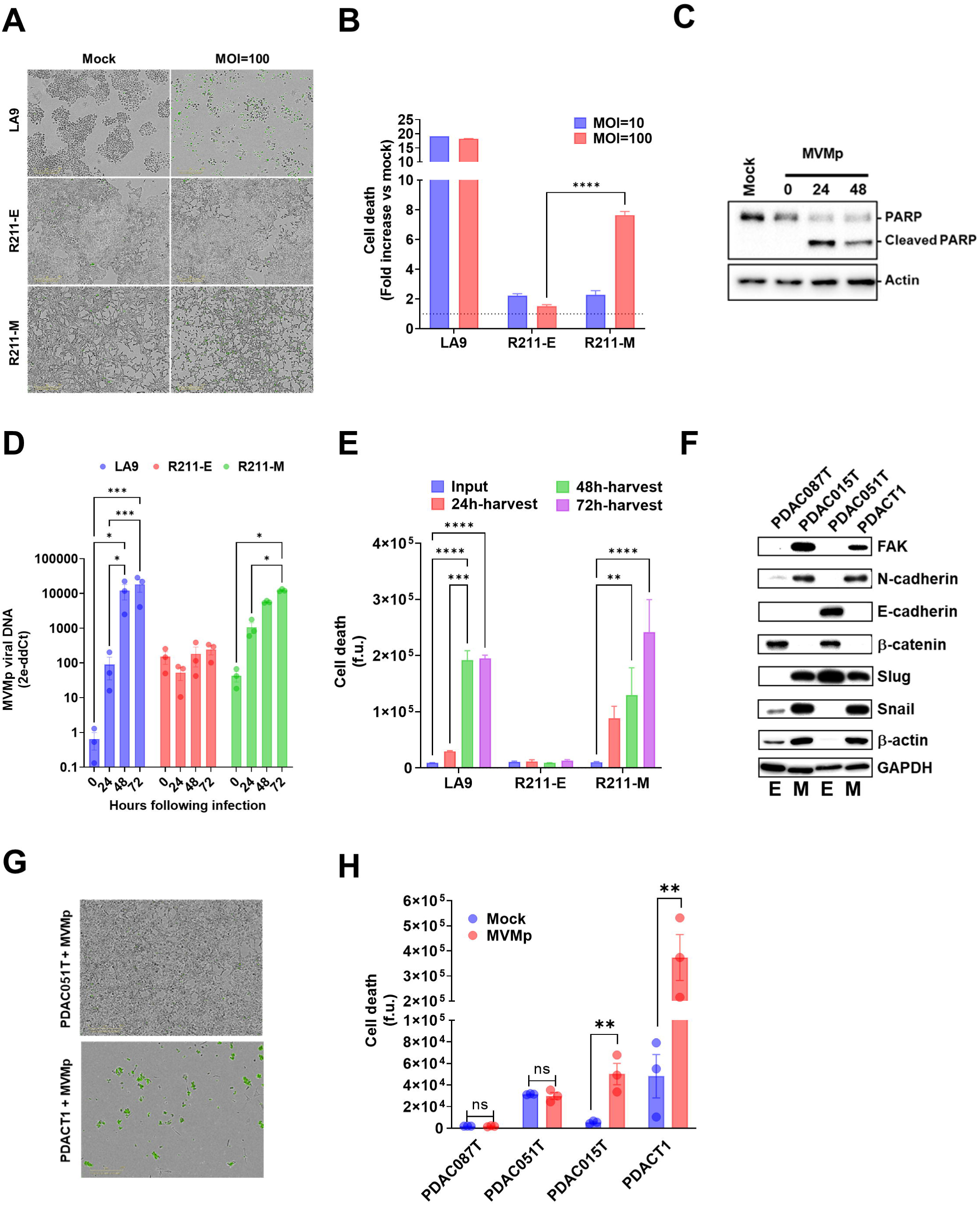
MVM is oncolytic in murine and human PDAC cells with mesenchymal phenotype. R211-M and R211-E cells were infected with MVMp at MOI=100. Control cells were incubated with infection medium. LA9 wells were used as positive control. Representative captions (**A**), real-time cell death analysis (**B**) and western blot for PARP cleavage of cells infected by MVMp or mock treated cells, up to 72 hours (**A**, **B**) or 48 hours (**C**) following infection. Results are mean ± s.e.m or representative of at least three independent experiments performed in duplicate. ****: p<0.001, 2-way Anova test. R211-M and R211-E cells were infected with MVMp at MOI=100, when LA9 were treated with 0.1M of MVMp. **D**. Cellular content in viral genomes was analyzed by qPCR for NS1 viral gene at the time indicated. Results are mean ± s.e.m of at three independent experiments performed in duplicate. *: p<0.05, ***::p<0.005, 2-way Anova test. **E**. Cell supernatant were collected at the time indicated and analyzed in real-time for oncolytic activity on LA9 reporter cells. Results are mean ± s.e.m of three independent experiments performed in duplicate. *: p<0.05, **: p<0.01, ***:p<0.005****:p<0.001, 2-way Anova test. **F**. Western blot analysis for the indicated proteins in human primary PDAC cells PDAC087T, PDAC015T, PDAC051T and PDACT1. Representative of two independent experiments. PDAC087T, PDAC015T, PDAC051T and PDACT1 were infected with MVMp at MOI=100 when control cultures were incubated with infection medium. E indicates epithelial, M stands for mesenchymal. Representative captions (**G**) and live cell death analysis (**H**) of human primary PDAC cells, 72 hours following infection by MVMp. Results are mean ± s.e.m of three independent experiments performed in duplicate. **: p<0.01, 2-way Anova test.

### MVMp specifically replicates in murine pancreatic cancer cells with mesenchymal phenotype

We next questioned whether murine PDAC cells may support viral replication. Thus, R211-M and R211-E cells were infected with MVMp at a MOI of 10. As control, LA9 cells were infected with MOI=0.1 of MVMp. Cells were collected at different time following infection and intracellular viral load was analyzed by qPCR. Figure 2D shows that LA9 cells replicate MVMp genome at very high level (more than 30000-fold increase as compared to viral input, 72h following infection, p<0.005, Figure 2D). On the contrary, R211-E cells failed to replicate MVMp genome (Figure 2D). Remarkably, R211-M cells significantly supported MVMp genome replication following infection, yet at lower level as compared to LA9 cells (130-fold increase as compared to viral input, 72h following infection, p<0.001, Figure 2D). These results demonstrate that murine PDAC cells with mesenchymal phenotype are permissive to MVMp infection.

### Viral progeny from MVMp-infected murine pancreatic cancer cells with mesenchymal phenotype is infectious

One key advantage of oncolytic virus like MVMp is that infected cells become viral factories allowing for several consecutive rounds of infection of neighboring cancer cells. Thus, we analyzed whether R211-M cells may produce and secrete infectious MVMp progeny following initial infection. R211-M and R211-E cells were infected with MVMp at a MOI of 10. As control, LA9 cells were infected with MOI=0.1 of MVMp. Following infection, cells were carefully washed with PBS and treated by proteinase K to remove viral input. Culture supernatants were collected at 0h, 24h, 48h and 72h following infection, and LA9 cells were used as reporter cells for oncolytic activity. Results shown Figure 2E indicate that cell supernatants from LA9 cells show lytic activity on reporter cultures, when supernatants collected from R211-E cells were ineffective. On the contrary, supernatant collected from R211-M cells infected by MVMp demonstrate a time-dependent increase lytic activity on reporter cells, indicating the presence of a fully oncolytic viral progeny (Figure 2E). Compared with cells with an epithelial phenotype, PDAC cells with a mesenchymal phenotype, that corresponds to the most aggressive PDAC phenotype, can be infected by MVMp to produce infectious viral progeny.

### MVMp specifically targets human primary pancreatic cancer cells with mesenchymal phenotype

To validate and extend these findings in patient-derived resources, we selected human PDAC primary cell cultures that presented with high expression of FAK, N-cadherin and Snail1 that are indicative of the “mesenchymal” phenotype (PDAC015T and PDACT1, Figure 2F) or high expression of E-cadherin and β-catenin, that signs the “epithelial” phenotype (PDAC087T, PDAC051T, Figure 2F). Primary cultures were infected with MVMp at MOI=100. PDAC087T and PDAC051T largely resisted infection (Figure 2F and 2G), while PDACT1 and, to a lesser extent PDAC051T, succumbed to MVMp infection (Figure 2F and 2G). These data corroborate MVMp tropism for mesenchymal-like cultures in human PDAC.

### MVMp shows antitumoral potential only in murine pancreatic cancer tumors with mesenchymal phenotype

To determine the antitumoral potential of MVMp *in vivo*, immune deficient mice with orthotopic R211-E or R211-M tumors were injected *i.v.* with 10^6^ or 10^7^ p.f.u. of MVMp, in two separate, 4-day distant injections (FigS3A). Control mice received PBS as placebo. Tumor growth monitoring by echography showed no impact of virotherapy on R211-E tumor growth (Fig2B). On the other hand, MVMp injection significantly reduced R211-M tumor growth in a dose dependent manner (−35%±3%, p<0.001, Figure 3A). Quantitative PCR for MVMp genome shows increased, yet not significant, level of viral load in R211-M tumors as compared to R211-E tumors (Figure 3B). To identify key tumor cell-intrinsic factors mediating MVMp antitumoral effect, we examined differentially expressed bulk transcriptomes from R211-M tumors receiving placebo or treated with 10e7 p.f.u. of the virus. Gene set enrichment analysis (GSEA) highlighted over expression of pathways related to cell senescence and TNF signaling (Figure 3D, Table S1) and inhibition of mitochondrial activity and DNA replication (Figure 3D, Table S2). In addition, these transcriptomes were enriched in genes and pathways related to apoptosis (Table S3). Consistent with this, western blot analysis revealed PARP cleavage in R211-M tumors treated by MVMp only, as compared to R211-E tumors (Figure 3E). Collectively, these data demonstrate the selectivity of MVMp for mesenchymal-like PDAC tumors *in vivo*.

**Figure 3.**
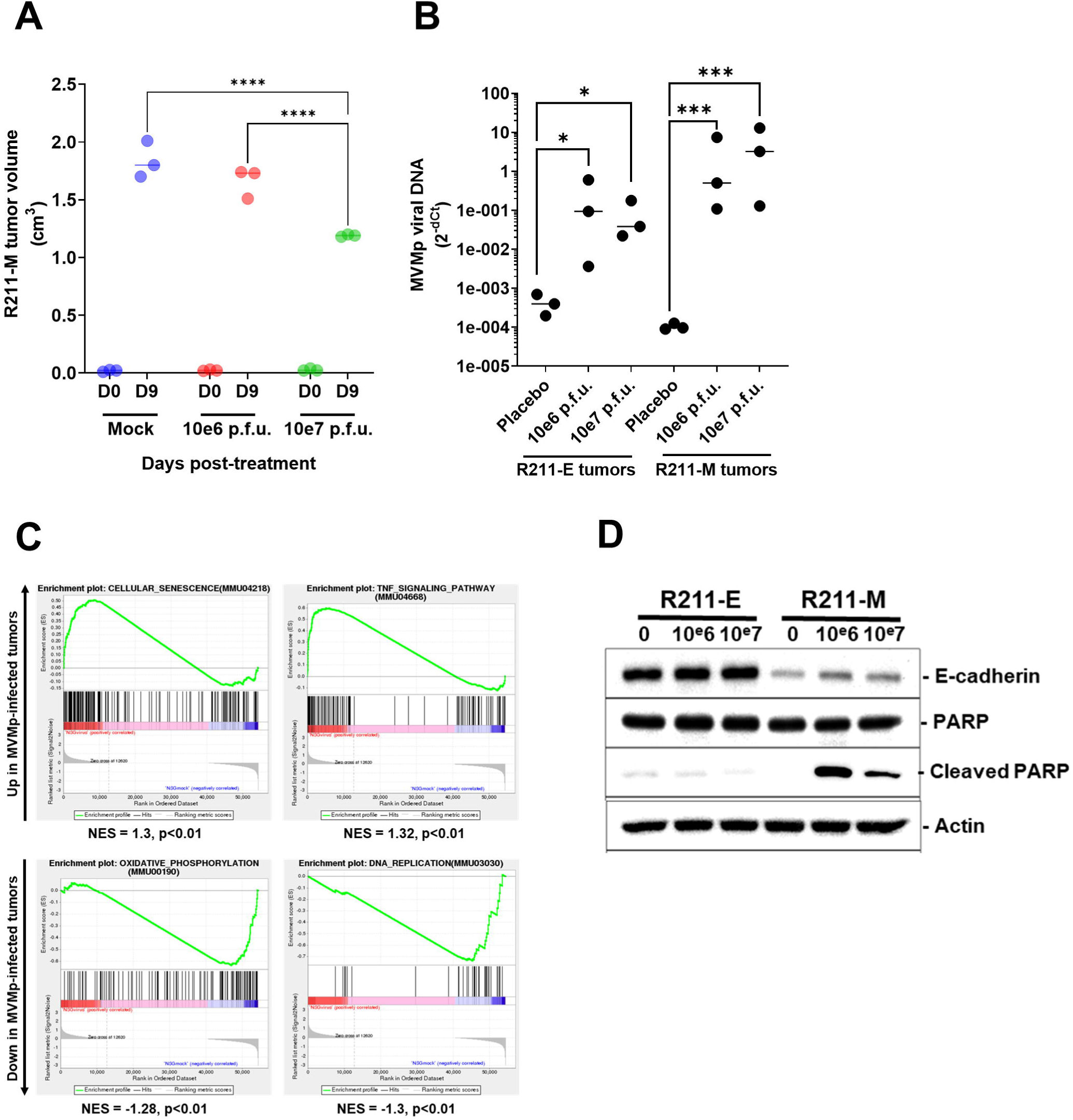
MVM is oncolytic in murine models of PDAC with mesenchymal phenotype. R211-M or R211-E tumors were induced in the pancreas of NSG mice as described in the Materials and Methods section. Eleven days later, when tumors reached 100 mm^3^ in size, mice were randomized and received two sequential *i.v.* injections of MVMp (10e6 or 10e7 p.f.u. for particle forming units), in two separate, 4-day distant injections (D0=day of the first injection). N=3 mice were used per group. **A.** R211-M tumor volume by echography, 9 days after starting treatment. Results are mean ± s.e.m or representative of three mice per group. ****: p<0.001, 2-way Anova test. At endpoint, tumors were sampled and tumor viral genome content was quantified by qPCR (**B**), and RNAseq followed by gene set enrichment analysis (GSEA, **C**) and Western blot for PARP cleavage (**D**) were performed. NES: normalized enrichment score. Results are mean ± s.e.m or representative of three mice per group. *: p<0.05, ***: p<0.005, 2-way Anova test.

### MVMp shows increased antitumoral efficacy in immune competent models of pancreatic cancer

OV replication within tumors is now largely assumed to cause direct lysis of cancer cells, leading to decrease of the bulk of the tumor. Along this line, preclinical and clinical data consistently suggest that OVs also utilize other anticancer mechanisms to combat tumors^19^. Previous work demonstrate that MVMp-infected glioma cells activate immune cells *in vitro*, and protect from tumor outgrowth *in vivo* in immunocompetent animals^10^. These data suggest that MVMp partners with the immune system to control tumor growth. However, the ability of MVMp to break immune tolerance in curative strategies has never been tested to date, especially in experimental models of PDAC. Accordingly, we generated orthotopic R211-M tumors in immune deficient (NSG) and immune competent (C56BL/6) mice models (FigS3C). Mice received two sequential *i.v.* injections of infra-therapeutic dose of MVMp (10e4 or 10e6 p.f.u.) as previously described. Control mice received PBS as placebo. At the lowest dose tested (10e4 p.f.u.) MVMp showed no effect on the growth R211-M tumors engrafted in immune deficient or immune competent mouse recipient (data not shown). At the highest dose tested (10e6 p.f.u), MVMp increased tumor cells death by apoptosis (FigS3D), was detected in tumors (FigS3E) but failed to inhibit R211-M experimental tumors growth when engrafted in immune deficient mice (Figure 4A and Figure 4C), consistent with our previous findings. Remarkably, infra-therapeutic MVMp injection significantly impaired the growth of very aggressive R211-M tumors in mice with intact immune system (−32%±5%, p<0.005, Figure 4B and Figure 4C). Mice survival was significantly extended when R211-M tumors were treated by the virus, as compared to control mice (15 days *vs* 11 days following initial virus injection, Log-rank Mantel-Cox test, p<0.01). We failed to detect tumor cell death induced by virotherapy (FigS3D), nor to retrieve viral genomes from infected R211-M tumors in C57BL/6 mice (FigS3E). On the other hand, antibodies against MVMp were readily detected by dot blot in the serum of infected mice (FigS3F). These findings strongly suggest that infected cells are rapidly sensed and cleared in immune competent animals.

**Figure 4.**
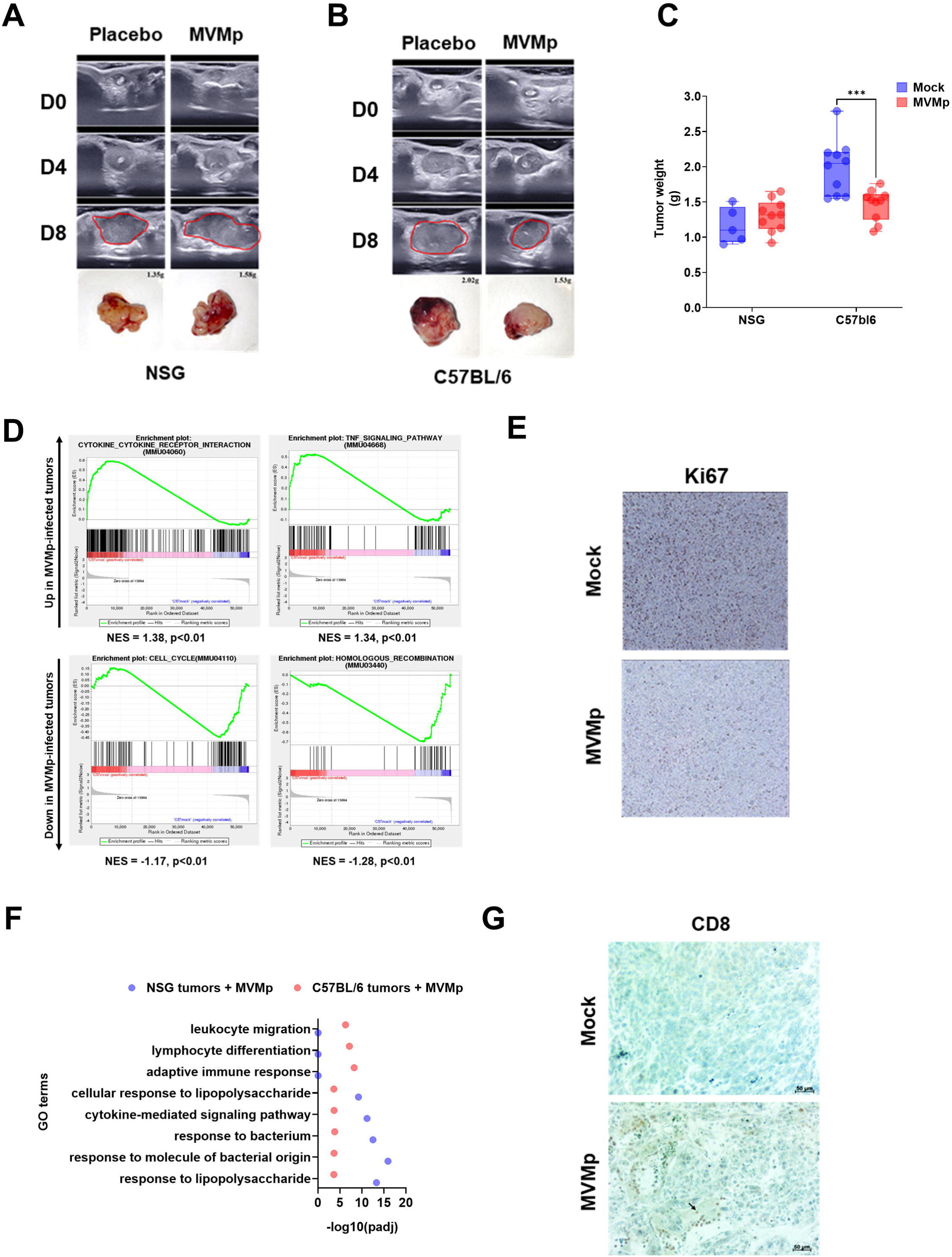
MVMp shows increased antitumoral efficacy in immune competent models of pancreatic cancer. R211-M tumors were induced in the pancreas of NSG mice or of C57BL/6 mice as described in the Materials and Methods section. When tumors reached 100 mm^3^ in size, mice were randomized and received two sequential *i.v.* injections of MVMp (10e4 or 10e6 p.f.u. for particle forming units), in two separate, 4-day distant injections (D0=day of the first injection). Eight to 10 mice were used per group. Representative echography captions and photographs with weight of R211-M tumors grown in NSG (**A**) or C57Bl6 (**B**) mice, up to 8 days following initial administration of MVMp. Tumors are circled in red. **C**. Tumor weight at endpoint. Results are mean ± s.e.m or representative of 8 to 10 mice per group. ***: p<0.005, 2-way Anova test. At endpoint, tumors were sampled and RNAseq followed by gene set enrichment analysis (GSEA, **D**), immunochemistry for Ki67 (**E**, R211-M tumors in C57Bl/6 mice only) gene ontology terms analysis (**F**) and immunochemistry for Ki67 (**G**, R211-M tumors in C57Bl/6 mice only) were performed in mice receiving 10e6 p.f.u. of the virus. NES: normalized enrichment score. Results are mean ± s.e.m or representative of 8 to 10 mice per group.

Next, we performed RNA sequencing on R211-M tumors engrafted in immune competent mice receiving placebo or 10e6 p.f.u. *i.v.* of the virus (Figure 3E). Tumor-treated transcriptomes were expectedly enriched in genes and pathways related to immune recruitment/function, while pathways indicative of cell proliferation were repressed (Figure 4D and Table S4). Tumor staining for Ki67 confirmed that tumor cell proliferation was significantly decreased following MVMp injection (−47.5%±5%, p<0.01, Figure 4E and FigS3G). To better understand the molecular basis for reduced tumor growth in the syngenic model of PDAC, we compared GO terms between MVMp-infected, R211-M tumors engrafted in NSG mice or in C57BL/6 mice. We found that pathways related to innate immune cells recruitment and function were similarly enriched in both models, while several pathways relative to adaptive immunity were only enriched in immune competent mice with tumors following MVMp administration (Figure 4F and Table S5). Immunochemistry analysis confirmed a dramatic increase in intratumoral CD8+ T cells in R211-M tumors engrafted in C57BL/6 mice and treated by MVMp, as compared to control tumors (Figure 4G). Collectively, these data demonstrate that MVMp virotherapy is more effective to inhibit the growth of autologous PDAC models, and increases effector T-cell infiltration in immune competent models of PDAC.

## DISCUSSION

The present study illustrates for the first time the high selectivity of MVMp oncolytic virus for a particular subtype of PDAC, a disease with no cure that highly resist to conventional therapies. MVMp was previously found to enter leading edge of migrating cells by endocytosis close to filopodial and focal adhesions areas, notably following induction of EMT^16^. Interestingly, two main molecular subtypes have been recently described in PDAC, including the classical/pancreatic progenitor subtype associated with a better prognosis, and the Basal-like/Squamous subtype associated with resistance to treatment and poorer clinical prognosis, including immune escape^20^. During this study, we built on the characterization of murine PDAC cells with either mesenchymal or epithelial phenotypes that highly resemble the basal-like and classical phenotypes found in patients with PDAC^2^. We demonstrate that infection by MVMp was cytotoxic and productive only in murine PDAC cells with mesenchymal phenotype, both *in vitro* and *in vivo,* as compared to cells with an epithelial phenotype that largely resisted virotherapy. To our knowledge, these data establish only the second example of a direct relationship between the mesenchymal phenotype of cancer cells with the increase in susceptibility to oncolytic infection. Indeed, Chen et al previously found that EMT enhances response to HSV-1 oncolytic virus therapy, probably due to enhanced cell surface expression of the viral receptor nectin-1^21^. Here, we went one step further as we identified a 5-protein classifier (FAK, N-cadherin, E-cadherin, β-catenin, Snail) that predicts response to MVMp virotherapy in murine PDAC models. We speculate that MVMp hijacks the uptake and lysosomal/proteasomal degradation of expired focal-adhesion molecules that are generated following physical tension of stress fibers, during the very dynamic process of mesenchymal cell migration, as previously described^16^. Considering non-migrating (epithelial) cells, where focal adhesion molecules remain very steadily attached and scarcely degraded, our data on viral replication indicate that the virus is either blocked outside the cell, trapped in endosomes, or brought back to the cell surface, unable to reach the nucleus and establish an infection. Electron microscopy experiments may help to better understand the mode of entry of MVMp in PDAC cells.

We also established an instructive role of this classifier in predicting MVMp lytic efficacy in patient-derived primary pancreatic tumor cells, albeit the “mesenchymal status” does not perfectly match the classical/basal-like molecular subtyping found in the literature^22^. Here, Slug expression seems not to be as important as in murine PDAC cells to predict MVMp efficacy. Further investigations are warranted to confirm the predictive value of the MVMp signature in more PDAC samples, but also in other tumor types with similar mesenchymal phenotype. As we and others documented that fine needle material can be sampled and analyzed following endoscopic exploration^23^, this finding opens for personalized, precision virotherapy for patients with PDAC.

Moreover, while R211-M cells successfully infected with the virus died by apoptosis, the non-infected cell population proliferated aggressively so that inhibition of cell proliferation was as best partial. The latter observation could be due to suboptimal viral replication and spread. Indeed, while detectable and infectious, viral progeny production by R211-M would be limiting to generate a second round of infection, as it may get diluted in the fast growing cell population and eventually degraded. Importantly, MVMp establishes replication at cellular DNA damage sites, that can be provoked by viral NS1 protein expression^9^, which provide replication and expression machinery^24^. Unfortunately, we failed to identify MVMp-induced DNA damage signaling in both R211-M and R211-E infected cells (Western blot for phosphorylated ATM and CHECK1, personal observation). One avenue to increase MVMp therapeutic efficacy will be to combine virotherapy with clinically relevant drugs such as gemcitabine or 5-Fluorouacil (5-FU), that induce DNA damage and that block cancer cells in G2/M or in S-phase, to create a situation that is more favorable to viral replication.

The present data not only build on a growing body of evidence implicating direct oncolysis of cancer cells, but also revealed novel insights into the immune potential of MVMp in experimental models of PDAC. This is of prime importance as pancreatic tumors are considered as immunologically dead and have not yet succumbed to immunotherapies^1^. To get one step closer to the clinical situation, we used quite unique experimental models where tumors were engrafted in the pancreas of recipient mice, and the virus administrated systemically. Despite this very challenging setting, MVMp significantly inhibited the progression of experimental R211-M tumors, when R211-E tumor growth was not impacted by virotherapy, further demonstrating the exquisite specificity of MVMp for the most aggressive subtype of PDAC. In immune compromised mice, MVMp was detected within tumors and associated with cell death by apoptosis, with evidence of expression of TNFα, that is both involved in viral sensing and type-1 antitumoral responses, inhibition of cancer cell proliferation, and engagement of innate immunity. Remarkably, infra therapeutic dosage of MVMp shows significant tumor growth inhibition in immune competent models, extended mice survival and promoted the massive infiltration of tumors by cytotoxic T cells. Our data confirm the general view that MVMp efficacy relies on the immune system; as previously found in experimental models of glioblastoma^10^. Additional experiments are needed to address the functional relevance of cytotoxic T cells in MVMp therapeutic efficacy in PDAC models, so as the level of exhaustion of intratumoral T cells, that can call for combination therapies of virotherapy with immune checkpoint blockers. Nevertheless, MVMp virotherapy comes with a cost. First, and as expected, mice produced antibodies against the virus following infection, a situation that may also occur in patients as MVMp is not a human pathogen. As active immune sensing of MVMp may blunt virotherapy efficacy (discussed below), this finding argues for a safe use of this biotherapy in immune competent patients. Second, our data suggest that infected cells are rapidly cleared by the adaptive immune system following infection, as no viral genome, nor apoptotic cells, were detected in tumors established in immune competent animals. This could result from the relatively modest oncolysis of tumor cells and of the probable expression of viral epitopes at the surface of infected cells. However, these findings may help revise the current, albeit highly challenged, *credo* for which “the antitumor activity of an oncolytic virus is proportional to the number of tumor cells it kills”. One could speculate that the presence of newly activated antiviral T cells may invigorate the pool of newly primed, cancer –specific cytotoxic effector T cells within the tumor bed (personal observation). Indeed, recent studies showed that pathogen-related CD4^+^ T cell memory populations can be re-engaged to support cytotoxic CD8^+^ T cells, converting a weak primary anti-tumor immune response into a stronger one. Along this line, immunization with the Newcastle Disease Virus (NDV) strongly increases both antiviral and antitumoral CD8^+^ T cells numbers, and tumor clearance before local treatment with NDV^25^. Consequently, vaccination strategies may improve the antitumoral activity of MVMp in syngenic models of PDAC.

In summary, the present study describes for the first time the oncolytic potential of MVMp in experimental models of PDAC. We show here that the highly selective tropism of MVMp may reveal a novel molecular vulnerability for patients with the most aggressive form of PDAC. Hence, we have identified a 5-gene classifier that may help select patients that would benefit the most from virotherapy. Last, we describe the immune potential of MVMp in syngenic tumor models, that can call for combination with immunotherapies. Leveraging such novel strategy could be of paramount importance to overcome therapeutic resistance in patients with PDAC.

## MATERIALS AND METHODS

### Cell culture

LA9 cells were a kind gift of Dr Nelly Panté (The University of British Columbia, Vancouver, BC, BC, Canada). R211 and R259 cells, that derive from spontaneous tumors generated in the KPC mice model (LSL-KrasG12D/+; LSL-Trp53R172H/+; Pdx-1-Cre), were obtained from D. Sauer (TUM, Munich, Ger). LA9, R211 and R259 cells were grown in DMEM supplemented with 4,5% glucose, 10% FBS and 1% Pen/Strep. Murine PDAC cells with mesenchymal phenotype (hereafter mentioned as R211-M or R259-M) were loosely attached to the culture dish and isolated following repeated pipetting in warm PBS. Murine cancer cells with epithelial phenotype (hereafter mentioned as R211-E or R259-E) strongly adhere to the plastic culture and were isolated following trypsinization. Pancreatic cancer patient-derived primary cells were obtained and cultured as previously described^26^. Cells were incubated at 37°C with 5% CO_2_.

### Virus production and purification

The initial MVMp stock was a kind gift of Nelly Panté (The University of British Columbia, Vancouver, BC, BC, Canada). All agreements for using MVMp were obtained from the French Ministry of Research, with authorization number DUO2333. Adherent LA9 cells were infected in high containment biological facility (Bl3, CRCT technological cluster) with MVMp at a MOI of 10^-2^, grown for 4 days at 37°C and 5% CO_2_, and harvested by scrapping in PBS. After a centrifugation at 2000*xg* for 5’, the cells were resuspended in TE (pH 8,7) and lysed through three cycles of freeze-and-thaw in liquid nitrogen. Further cell lysis was obtained by direct sonication at 10W (4 times 3’’) on ice, and cell debris were pelleted by centrifugation at 15000*xg* for 1h. The virus was then precipitated by addition of CaCl2 (25mM final), incubated on ice for 30min, then spun at 10000*xg* for 15min at 4°C and resuspended in DNAse I (50U/ml) to eliminate free viral DNA. After a 20min incubation at 37°C, the precipitation step was repeated and the pellet of virus was resuspended in TE. The process of sonication was then repeated as described above, and cell debris were eliminated by centrifugation at 15000*xg* for 1h at 4°C. The virus was finally loaded on Sucrose/CsCl gradients for a 24h centrifugation at 100000*xg* and 4°C. The virus was extracted and dialyzed 3 times overnight against TE, and then filtered through a 0,2μm PES syringe filter.

### Plaque assay

Cells were seeded in 6cm dishes at 6.10^3^ (R211) or 2.10^5^ (LA9) cells/ml (5ml/dish) in complete DMEM (4.5% glucose, 10% FBS and 1% Pen/Strep) and grown at 37°C and 5% CO_2_ for 24h. After a brief wash in PBS, the cells were incubated for 1h at 37°C with 10X dilutions (in Infection Medium: DMEM 1% FBS and 1mM Hepes) of purified MVMp, starting at a MOI of 100. As a control (MOCK), cells were incubated in infection medium alone. The supernatant was then removed and 7ml of overlay medium (MEM containing 0.5% glucose, 5% FBS, 0I2% gentamycin, 1% triptose phosphate and 0,75% low melting-point agarose) were added to each dish and allowed to solidify for 30min at RT. The cells were then grown at 37°C and 5% CO_2_ for 4 (R211) or 5 (LA9) days, then fixed in 4% formaldehyde for 30min and stained with 0.3% methylene blue for 30min.

### Virus infections and analysis

Cells were seeded in 6-well plates at 5.10^3^ (R211) or 2.10^4^ (LA9) cells/ml (2ml/well) in complete DMEM and grown at 37°C and 5% CO_2_ for 24h. After a brief wash in PBS, the cells were incubated for 1h at 37°C with purified MVMp diluted in infection medium, at the indicated MOIs. As a control (MOCK), cells were incubated in infection medium alone. The supernatant was then removed, 2ml of complete DMEM containing 10nM Cytotox-Green (Sartorius) were added to each well, and the cells were grown at 37°C and 5% CO_2_ in an Incucyte Zoom (Sartorius). Cell death was monitored hourly.

### MVMp secretion analysis

Cells were seeded in 10cm dishes at 3.10^4^ (R211) or 7.10^4^ (LA9) cells/ml in complete DMEM and grown at 37°C and 5% CO_2_ for 24h. After a brief wash in PBS, the cells were incubated for 1h at 37°C with purified MVMp diluted in infectious medium, at a MOI of 10 (R211) or 0.1 (LA9). As a control (MOCK), cells were incubated in infectious medium only. The supernatant was then removed and the cells were incubated for 30min at RT in PBS containing 0.1mg/ml Proteinase K (1ml/plate) for 30min at RT. The cells were subsequently collected, spun at 2000*xg* for 5’ and washed in 1ml PBS. After another centrifugation at 2000*xg* for 5’, the cells were resuspended in complete DMEM and finally reseeded in 10cm dishes and grown at 37°C and 5% CO_2_. Culture supernatants were collected at 0h, 24h, 48h and 72h post-reseeding for Incucyte-based analysis of MVMp secretion using LA9 as a reporter cell. For this latter experiment, LA9 cells were seeded in 6-well plates at 2.10^4^ cells/ml in complete DMEM (2ml/well) and grown at 37°C and 5% CO_2_ for 24h. After a brief wash in PBS, the cells were incubated for 1h at 37°C with 1μl of each collected supernatant diluted 1:100 in infection medium. The supernatants were then aspirated and 2ml of complete DMEM containing 10nM Cytotox-Green (Sartorius) were added to each well. The cells were grown at 37°C and 5% CO_2_ in an Incucyte Zoom (Sartorius) where cell death was monitored hourly.

### Western and dot blots

For western blot analysis, cells were scrapped in PBS, spun at 2000*xg* for 5min at 4°C, resuspended in Radioimmunoprecipitation assay buffer (RIPA, Bio Basic, CA) lysis buffer containing protease inhibitors (Sigma), and incubated on ice for 30min. After a 10min centrifugation at 15000*xg* and 4°C, supernatants were collected and protein concentration determined by Bradford assay (Bio-Rad). For the preparation of tumor samples, 2-3mm cubes of tumors were mixed with 1mm glass beads and 300μl of ice-cold RIPA lysis buffer containing protease inhibitors, then lysed using Precellys tissue homogenizer (Bertin) through 4 cycles of 10sec at 800rpm. After a 10min centrifugation at 15000*xg* and 4°C, supernatants were collected and protein concentration determined by Bradford assay (Bio-Rad). Depending on the proteins of interest, 8% or 14% SDS-PAGE gels were loaded with 40μg or 80μg of total proteins (diluted in 5X sample buffer and boiled for 5min at 96°C) and run for 2h at 140V. After semi-dry transfer of the proteins onto nitrocellulose membranes (Bio-Rad), blocking was performed in PBS 5% skim milk for 1h at RT. Primary antibodies (Cell signaling technology) were incubated overnight at 4°C as per recommended by the manufacturer, and HRP-coupled secondary antibodies (Bio-Rad) for 2h at RT. For Dot blot studies, 1μl drops containing 10^4^ pfu of purified MVMp were dried onto a nitrocellulose membrane. After blocking in PBS 5% skim milk for 1h at RT, membranes were incubated overnight at 4°C with serums from MVMp or MOCK treated mice diluted 1:1000 in PBS. After 3 washes in PBS, membranes were incubated with HRP-coupled anti-mouse secondary antibodies (1/10^4^) for 2h at RT, and then washed again 3 times before revelation by electrochemoluminescence (Bio-Rad) using the Chemi-doc XRS+ (Bio-Rad).

### Viral genome quantification

Cells were seeded in 10cm dishes at 3.10^4^ (R211) or 7.10^4^ (LA9) cells/ml (10ml/dish) in complete DMEM and grown at 37°C and 5% CO_2_ for 24h. After a brief wash in PBS, the cells were incubated for 1h at 37°C with purified MVMp diluted in infectious medium, at MOIs of 10 (R211) or 0.1 (LA9). The cells were subsequently grown at 37°C and 5% CO_2_ and harvested by scraping in 1ml PBS at 0h, 24h, 48h or 72h post-infection, then spun at 2000*xg* for 5min. Cell pellets were resuspended in ice cold TE (pH 8.7), and lysed through three cycles of freeze-and-thaw in liquid nitrogen. For the preparation of tumor samples, 2-3mm^3^ of tumors were mixed with 1mm glass beads and 300μl of ice-cold TE, then lysed using a Precellys tissue homogenizer (Bertin) through 4 cycles of 10sec at 800rpm. After a 30min centrifugation at 15000*xg* and 4°C, 10μl of supernatant was mixed with 30μl of 1M NaOH (diluted in TE) and immediately incubated at 56°C for 30min to release viral genomes. The reaction was finally stopped by the addition of 960μl of 40mM HCl. For qPCR analysis, 1μl of sample was mixed in 20μl final of SoFast-Green reaction mix containing 10nM of forward (CCGTCTTAAGTTTGATTTT) and reverse (AGAGGTGGACCAACTCGGTA) primers for MVMp NS1 gene amplification using the StepOnePlus real-time PCR system (Thermo Fisher Scientific). Primers targeting the 18S gene were used for normalization (forward: AAACGGCTACCACCATCCG, reverse: CCTCCAATGGATCCTCGTTA). Relative quantity of viral DNA was calculated by the comparative threshold cycle (CT) method as 2^−ΔCT^.

### Animal experimentation

Experimental procedures performed on mice were approved by the ethical committee of INSERM CREFRE US006 animal facility and authorized by the French Ministry of Research: APAFIS#3600-2015121608386111v3. Six to 8 weeks- old NSG or C57BL/6 mice were surgically grafted orthotopically with 5.10^4^ R211-M or R211-E cells diluted in 50μl total PBS per animal. One week later, tumor formation was measured non-invasively by echography using the Aixplorer imaging device (Supersonic Imagine). Four days later, echography was performed and groups of mice with comparable average tumor sizes were made. Randomized mice were injected intravenously with 50μl of purified MVMp (diluted 1:10 in PBS) in high containment animal facility (A3 level). As placebo, mice received a similar volume of TE diluted 1:10 in PBS. Tumor volume was measured at the indicate days using the Aixplorer imaging device. For dot blots experiments, blood was sampled and spun at 2000*xg* for 5min, and serum was frozen immediately.

### Immunohistochemistry

Samples were incubated at 60°C for 1h, then incubated 3x in xylene for 5min. Samples were then rehydrated by incubation in 100% ethanol (3x 5min), 90% ethanol (2X 5min), 70% ethanol for 2min, 50% ethanol for 5min, then 2X in dH_2_O for 5min each. After a 10min permeabilization step in 0.1% Triton X-100 in PBS, samples were washed 3X in milliQ dH_2_O. For antigen retrieval, samples were autoclaved at 121°C for 12min in citrate buffer (20mM citric acid, 1mM sodium citrate), allowed to cool for 1h and incubated 3X 5min in PBST. For sample staining, slides were placed in humid chambers, blocked in Protein block (DAKO) for 10min at RT, then incubated overnight at 4°C with KI67 primary antibody (cell signaling technology) diluted 1:400 or CD8a primary antibody (Biolegend) diluted 1:200 in Antibody diluent signal stain (Dako). After 3 wash of 5min in PBS-Tween, peroxidase activity was inhibited by incubation of the samples in PBST-H_2_O_2_ (3%) for 15min at RT. The samples were then washed twice in PBS-Tween, once in PBS-Tween 1% BSA (5min each), and subsequently incubated with SignalStain Boost IHC detection reagent (cell signaling technology) for 45min at RT, in humid chambers. The samples were then washed 3X in PBS 1% BSA, once in PBS-Tween, incubated in AEC (Dako) for 5min, and washed twice for 5min in PBST. For counterstaining, samples were incubated for 2min in hematoxylin (Merck) and washed for 5min in dH_2_O, then mounted in Glycergel (Dako) and analyzed using an Axiolab A1 inverted microscope (Zeiss).

### RNA isolation and gene expression analysis

Tumors were sampled and minced and total RNA was extracted from tumors with RNeasy kit (Qiagen) according to the manufacturer’s instructions; quality and quantity were measured on a Nanodrop system (ThermoFisher Scientific). RNAseq analysis (polyA capture) was performed using 1µg of total RNA and subcontracted to Novogene. Briefly, RNA degradation and contamination was monitored on 1% agarose gels. RNA purity was checked using the NanoPhotometer® spectrophotometer (IMPLEN, CA, USA). RNA integrity and quantitation were assessed using the RNA Nano 6000 Assay Kit of the Bioanalyzer 2100 system (Agilent Technologies, CA, USA). A total amount of 1 μg RNA per sample was used as input material for the RNA sample preparations. Sequencing libraries were generated using NEBNext® UltraTM RNA. Library Prep Kit for Illumina® (NEB, USA) following manufacturer’s recommendations and index codes were added to attribute sequences to each sample. Briefly, mRNA was purified from total RNA using poly-T oligo-attached magnetic beads. Fragmentation was carried out using divalent cations under elevated temperature in NEBNext First Strand Synthesis Reaction Buffer (5X). First strand cDNA was synthesized using random hexamer primer and M-MuLV Reverse Transcriptase (RNase H-). Second strand cDNA synthesis was subsequently performed using DNA Polymerase I and RNase H. Remaining overhangs were converted into blunt ends via exonuclease/polymerase activities. After adenylation of 3’ ends of DNA fragments, NEBNext Adaptor with hairpin loop structure were ligated to prepare for hybridization. In order to select cDNA fragments of preferentially 150∼200 bp in length, the library fragments were purified with AMPure XP system (Beckman Coulter, Beverly, USA). Then 3 μl USER Enzyme (NEB, USA) was used with size-selected, adaptorligated cDNA at 37 °C for 15 min followed by 5 min at 95 °C before PCR. Then PCR was performed with Phusion High-Fidelity DNA polymerase, Universal PCR primers and Index (X) Primer. At last, PCR products were purified (AMPure XP system) and library quality was assessed on the Agilent Bioanalyzer 2100 system. The clustering of the index-coded samples was performed on a cBot Cluster Generation System using PE Cluster Kit cBot-HS (Illumina) according to the manufacturer’s instructions. After cluster generation, the library preparations were sequenced on an Illumina platform and paired-end reads were generated.

Raw data (raw reads) of FASTQ format were firstly processed through fastp. In this step, clean data (clean reads) were obtained by removing reads containing adapter and poly-N sequences and reads with low quality from raw data. At the same time, Q20, Q30 and GC content of the clean data were calculated. All the downstream analyses were based on the clean data with high quality. Reference genome and gene model annotation files were downloaded from genome website browser (NCBI/UCSC/Ensembl) directly. Paired-end clean reads were mapped to the reference genome using HISAT2 software. HISAT2 uses a large set of small GFM indexes that collectively cover the whole genome. These small indexes (called local indexes), combined with several alignment strategies, enable rapid and accurate alignment of sequencing reads. Differential expression analysis between two conditions/groups (three biological replicates per condition) was performed using DESeq2 R package. DESeq2 provides statistical routines for determining differential expression in digital gene expression data using a model based on the negative binomial distribution. The resulting P values were adjusted using the Benjamini and Hochberg’s approach for controlling the False Discovery Rate (FDR). Genes with an adjusted P value < 0.05 found by DESeq2 were assigned as differentially expressed. Gene Ontology (GO) enrichment analysis of differentially expressed genes was implemented by the clusterProfiler R package, in which gene length bias was corrected. GO terms with corrected Pvalue less than 0.05 were considered significantly enriched by differential expressed genes. We used clusterProfiler R package to test the statistical enrichment of differential expression genes in KEGG pathways.

### Data Availability Statement

Raw transcriptomic data supporting the findings of this study are available from the corresponding author upon reasonable request.

### Ethics reporting

Experimental procedures performed on mice were approved by the ethical committee of INSERM CREFRE US006 animal facility and authorized by the French Ministry of Research: APAFIS#3600-2015121608386111v3

### Statistical analysis

Grouped analysis were performed using two-way Anova tests with multiple comparisons using GraphPad Prism 9 software. Survival analysis were performed using a Log-rank (Mantel-Cox) test. Values are presented as \**p*L<L0.05, \*\**p*L<L0.01 and \*\*\**p*L<L0.001 and **** *p*L<L0.0001. Error bars are s.e.m. unless otherwise stated. The experiments were performed with a sample size *n* greater than or equal to three replicates, and the results from representative experiments were confirmed in at least two independent experiment repeats. When monitoring tumor growth, the investigators were blinded to the group allocation but were aware of group allocation when assessing the outcome. No data were excluded from the analyses.

## Acknowledgments

The authors are grateful to Dr Nelly Panté (The University of British Columbia, Vancouver, BC, BC, Canada) for the kind gift of LA9 cells and MVMp virus. The authors thank the financial support of “Fondation ARC” (PG fellowship, 2016) and of Inserm Transfert. The authors thank Ms Emilie Martin and Catherine Zanibellato for technical assistance. We are grateful to the Non-Invasive Exploration service (UMS006/CREFRE, Anexplo Platform, Toulouse) for giving us the access to the Aixplorer, that was acquired with financial support from ITMO Cancer AVIESAN (Alliance Nationale pour les Sciences de la Vie et de la Santé, National Alliance for Life Sciences & Health) within the framework of the Cancer Plan. The authors are grateful to Loïc Van Den Berghe and Tiphaine Fraineau(Vectorology facility, Technology Cluster of the Cancer Research Center of Toulouse, INSERM- UMR1037) for their technical assistance.

## Author contributions

Conceptualization: Pierre Garcin, Pierre Cordelier, Neslon Dusetti, Hubert Lulka, Louis Buscail. Investigations : Pierre Garcin, Hubert Lulka, Pierre Cordelier. Data analysis: Pierre Garcin, Margaux Vienne, Nelson Dusetti, Louis Buscail, Pierre Cordelier. Supervision : Pierre Cordelier. Writing – original draft: Pierre Garcin and Pierre Cordelier. writing – review & editing: Pierre Cordelier and Margaux Vienne. Funding acquisition : Pierre Cordelier.

---

**Figure S1.**
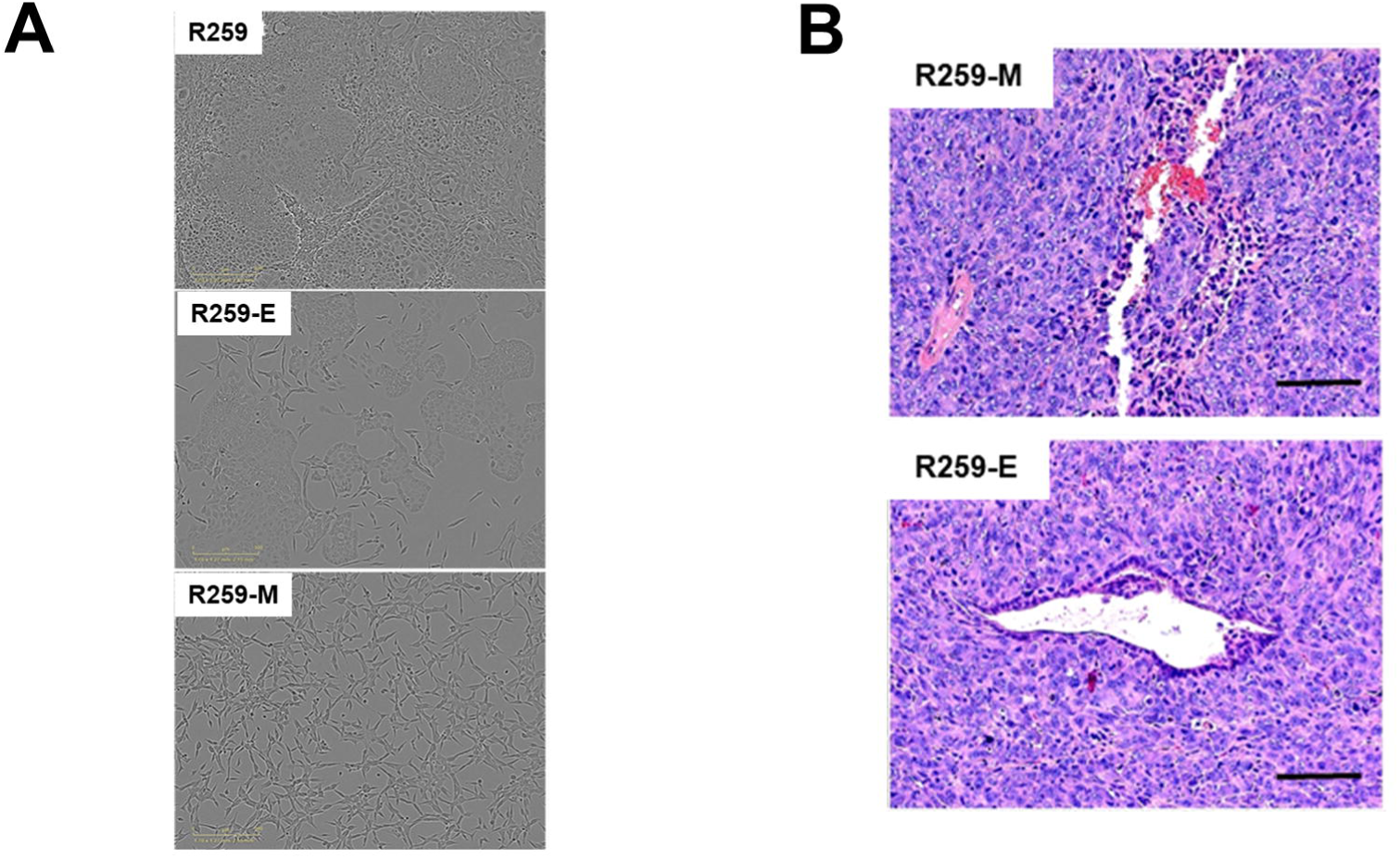
R259-E and R259-M cells were isolated by differential adherence to plastic as mentioned in the Materials and Methods section. **A.** Representative captions of long term culture of R259 cells (top) and of isolated R211-E (middle) and R211-M (bottom) cells. **C.** hematoxylin and eosin (HE) staining of orthotopic R259-E and R259-M tumors engrafted in nude mice at endpoint. Representative of n=3 mice per group.

**Figure S2.**
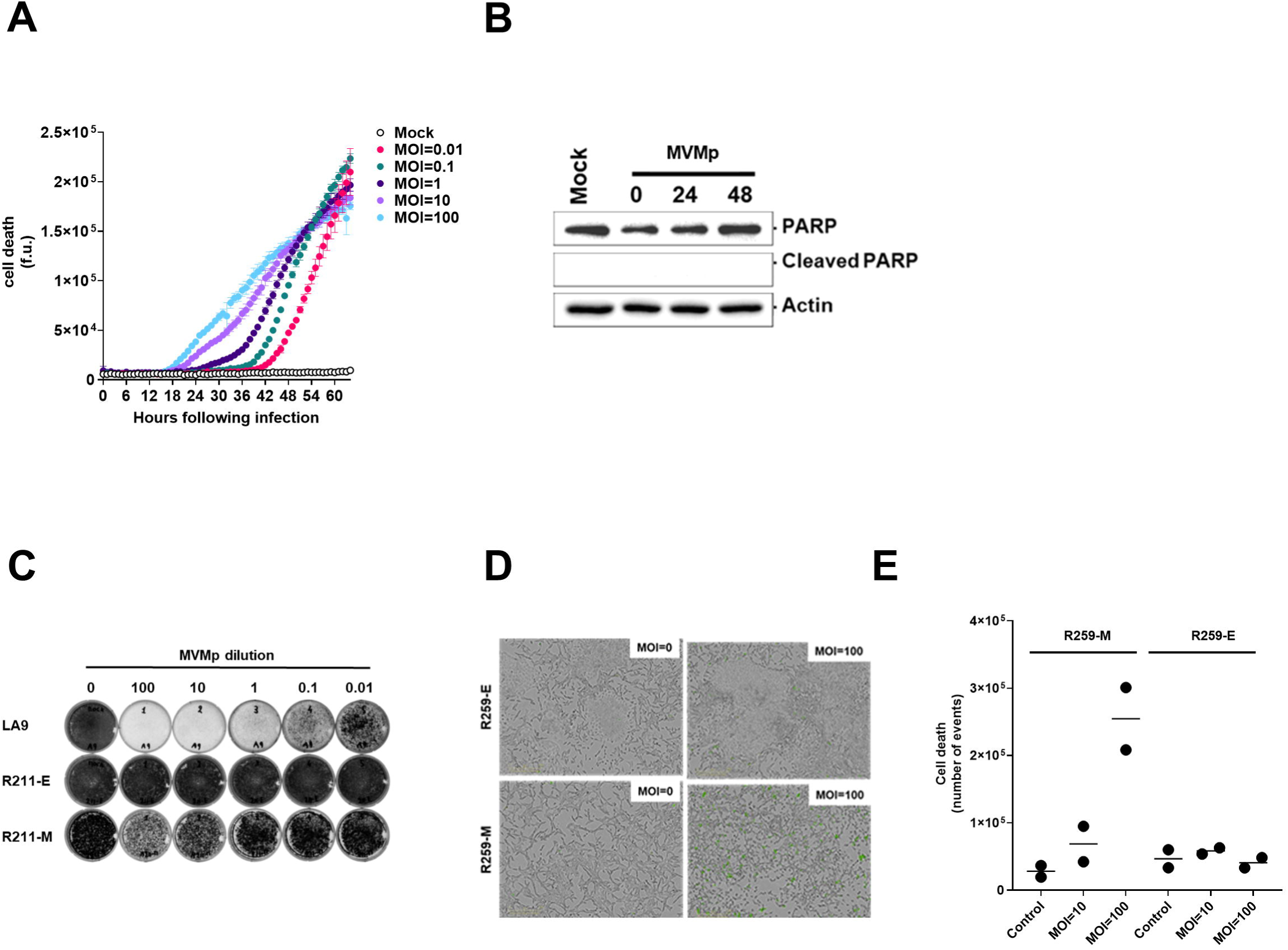
**A**. Real-time, dose-dependent analysis of MVMp- induced cell death in duction in LA9 cells. Results are mean ± s.e.m of three independent experiments performed in duplicate. **B**. Western blot for PARP cleavage in R259-E cells infected with MVMp at MOI=100, up to 48 hours following infection. Result is representative of two independent experiments. **C**. Plaque assay for MVMp at the indicated dilution in R211-M and R211-E cells. LA9 cells were used as positive control. Result is representative of three independent experiments. R259-M and R259-E cells were infected with MVMp at MOI=100. Control cells were incubated with infection medium. Representative captions (**D**) and real-time cell death analysis (**E**) of cells infected by MVMp or mock treated cells, 72 hours following infection. Results are mean of two independent experiments performed in duplicate.

**FigS3.**
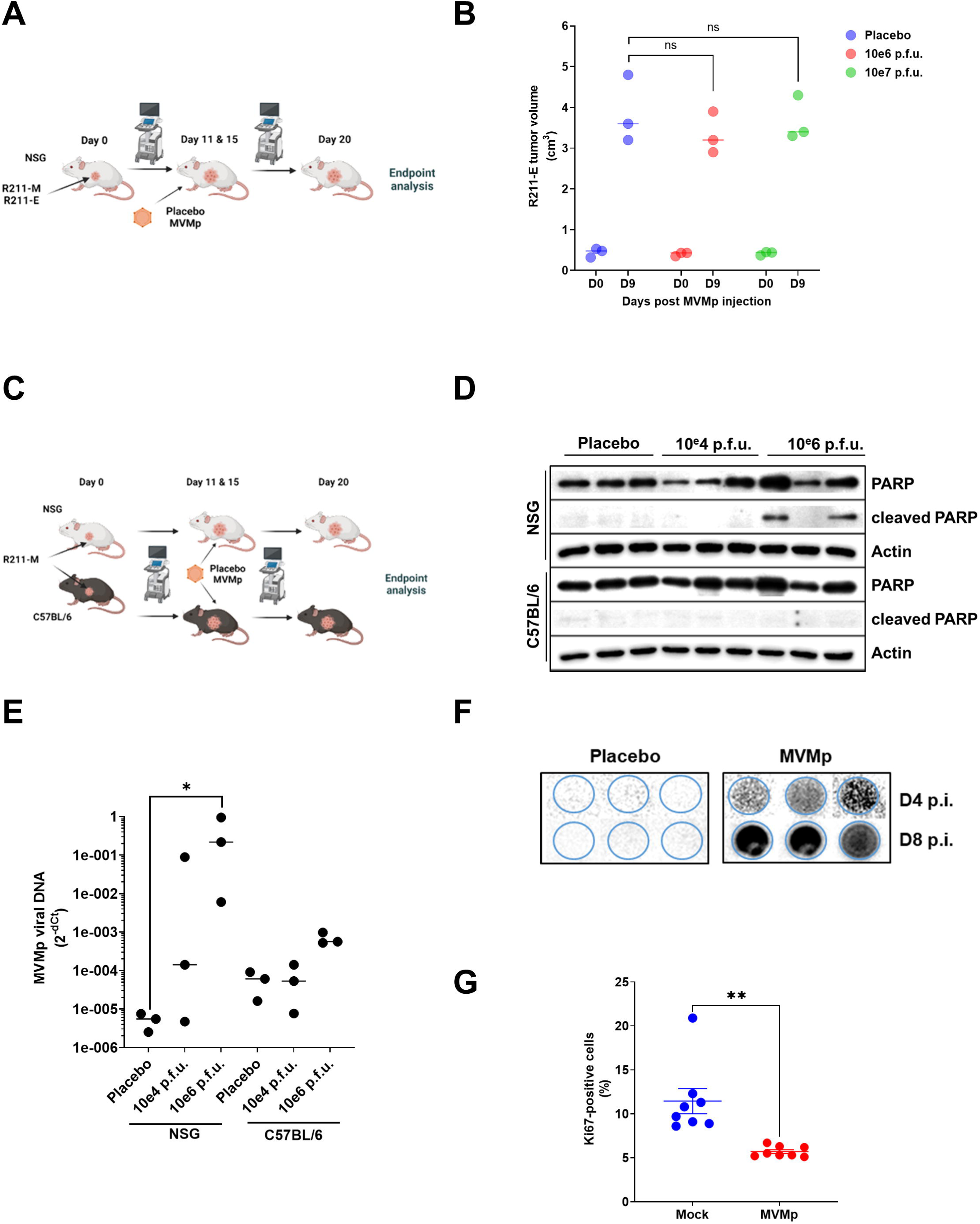
**A**. Experimental setting for the analysis of MVMp antitumoral efficacy on R211-M and R211-E tumors engrafted in NSG mice. **B**. R211-E tumors were induced in the pancreas of NSG mice as described in the Materials and Methods section. Eleven days later, when tumors reached 100 mm^3^ in size, mice were randomized and received two sequential *i.v.* injections of MVMp (10e6 or 10e7 p.f.u. for particle forming units), in two separate, 4-day distant injections (D0=day of the first injection). N=3 mice were used per group. R211-E tumor volume was assessed by echography. **C**. Experimental setting for the analysis of MVMp antitumoral efficacy on R211-M tumors engrafted in NSG or C57BL/6 mice. R211-M tumors were induced in the pancreas of NSG mice or of C57BL/6 mice as described in the Materials and Methods section. When tumors reached 100 mm^3^ in size, mice were randomized and received two sequential *i.v.* injections of MVMp (10e4 or 10e6 p.f.u. for particle forming units), in two separate, 4-days distant injections (D0=day of the first injection). Eight to 10 mice were used per group. Western blot for PARP cleavage (**D**) and viral genome quantification by qPCR (**E**) were performed at endpoint. Results are mean or representative of three independent experiments. *: p<0.05. 2-way Anova test **F**. Mice serum receiving placebo or 10e6 p.f.u. wasa assayed for the presence of MVMp antibodies at the indicated time. N=3 mice were used per group. At endpoint, R211-M tumors grown in C57BL/6 mice receiving or not 10e6 p.f.u. of MVMp were assayed for Ki67 staining. **G**. Results are mean ± s.e.m of the percentage of Ki67 positive in 2-3 fields from 3 different tumors. **: p<0.05. Unpaired Student T-test.

